# ‘Merging of ventral fibers at adhesions drives the remodeling of cellular contractile systems in fibroblasts.’

**DOI:** 10.1101/2022.03.16.484620

**Authors:** Shwetha Narasimhan, William R. Holmes, Irina Kaverina

**Affiliations:** Department of Cell and Developmental Biology, Vanderbilt University; Department of Physics and Astronomy, Vanderbilt University

**Keywords:** Actin Stress Fiber, Ventral Stress Fiber, Focal Adhesion, Cytoskeleton, Stress fiber dynamics

## Abstract

Ventral stress fibers (VSFs) are contractile actin fibers present in the ventral plane of the cell and existing in a dynamic attachment with cell-matrix focal adhesions. VSFs are critical in cellular mechanobiological functions such as traction force production, cell polarization, and migration. VSF within their intracellular network vary from short, thinner fibers that are randomly oriented to long, thick fibers that span along the whole long axis of a cell. *De novo* VSF formation was shown to occur by condensation from the cortical actin mesh or by crosslinking of other stress fiber subtypes (dorsal stress fibers and transverse arcs) at the cell front. However, formation of long VSFs that extend across the whole cell axis is not well understood. Here, we report a novel phenomenon of VSF merging in migratory fibroblast cells, which is guided by mechanical force balance and contributes to VSF alignment along the long cell axis. The mechanism of VSF merging involves two steps: connection of two ventral fibers by an emerging myosin II bridge at an intervening adhesion and intervening adhesion dissolution to form a cohesive, contractile VSF. Our data indicate that these two steps are interdependent, since under conditions where adhesion disassembly is slowed, formation of the myosin bridge is slowed as well. Cellular data and computational modeling show that the angle of contact between merging fibers decides successful merging, with angles closer to 180 yielding merging events and shallower angles leading to merge failure. Our data and modeling further show that merging increases the share of uniformly aligned long VSFs, which would contribute to directional traction force production. Thus, we thoroughly characterize merging as process for dynamic reorganization of VSFs in steady state, investigating the steps and variants of the process as well as its functional significance in migratory cells.

## Introduction

The actin cytoskeleton in a cell plays an integral role in cell polarization, migration, and force production [1]. Filamentous actin structures in migrating cells include branched and interlocked networks found in lamellipodium and the cortical mesh, parallel actin bundles in filopodia and antiparallel actin bundles in stress fibers [2]. Actin stress fibers are higher order structures formed by crosslinked bundles of 10-30 actin filaments [3]. Actin stress fibers and focal adhesions are closely linked in terms of structure and dynamics [4,5,6]. Actin polymerization initiates in nascent focal contacts [7], while further maturation of focal adhesions depends on both the tension applied and the structural template offered by the growing actin filament [8].

Stress fibers in a migrating mesenchymal cell can be divided into three main subtypes based on their location and attachment to focal adhesions – dorsal stress fibers, transverse arcs and Ventral Stress Fibers (VSFs), as depicted in ***Fig 1A*** [9,10,11,12]. Dorsal stress fibers are non-contractile fibers that are crosslinked by alpha-actinin. They are attached to the substrate through a single focal adhesion at their distal end and rise from the bottom to the top of the cell [13,14]. Transverse arcs are curved, contractile, α-actinin- and myosin II-associated actin bundles that form an interconnected network with dorsal stress fibers, rising to the upper planes of the cell. Transverse arcs are not attached to focal adhesions, and their contractile forces are transduced to the substrate by dorsal stress fibers and their associated adhesions [13,15,16].

**Figure 1:**
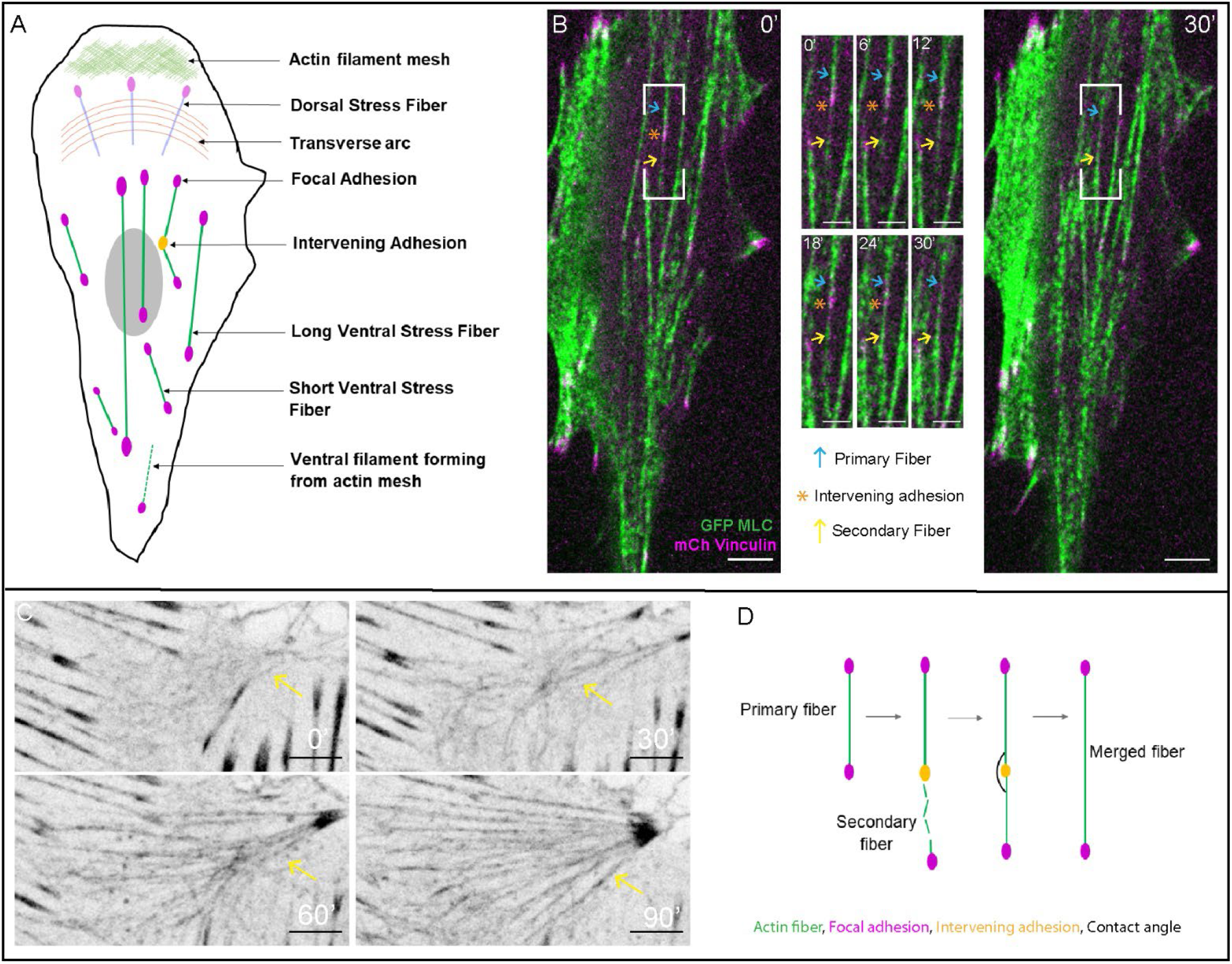
Ventral stress fibers remodel by merging at intervening focal adhesions **A)** Schematic of ventral view of a migrating cell with different actin stress fibers marked. The green fibers denote ventral stress fibers (VSFs) at the bottom plane attached to magenta focal adhesions on either end. VSFs are of many lengths and variably oriented along the cell’s main axis. Two VSFs are shown merging with an intervening adhesion between. Formation of actin fiber by coalescence from the actin mesh is marked. Stress fiber subtypes in upper z planes such as dorsal stress fibers and transverse arcs are also shown in faded colors. **B)** Ventral plane timelapse of MRC5 cell plated on 1 μg/ml Fibronectin substrate expressing GFP-Myosin Light Chain (MLC, green) and mCherry Vinculin (magenta) showing a merging event. Boxes on cell images at 0 min and 30 min note position of the merge event in the insets. Insets show merge event where a newly formed secondary ventral fiber (yellow arrow) is attached to the primary ventral fiber (Blue arrow) at the intervening adhesion (orange asterisk). As the intervening adhesion dissolves, the merging fibers are joined into a single merged ventral fiber by 30 min. Scale bar= 10 μm (all). See Suppl Video 1. **C)** Ventral plane timelapse of MRC5 cell plated on 10 μg/ml Fibronectin substrate expressing GFP-Utrophin showing secondary fiber formation by actin mesh condensation. The disordered actin mesh in 0 min condenses spontaneously into filaments. Scale bar= 10 μm See Suppl Video 2. **D)** Schematic of merging showing a pre-existing primary fiber attached to two adhesions. Secondary fiber forms and attaches to an end adhesion, which becomes the intervening adhesion. As the intervening adhesion disassembles, the two fibers are merged into one. Black arc denotes contact angle of the merging fibers.

Ventral stress fibers (VSFs) are arguably the most significant actin assemblies in a migrating mesenchymal cell. They are prominent, discrete contractile fibers in the ventral plane of the cell, consisting of actin filaments crosslinked by alpha-actinin and associated with myosin II stacks. They are attached to the substrate through a focal adhesion at each end. VSFs exist in many lengths and various angles with respect to the main cell axis, with longer VSFs more aligned towards the major axis of the cell. They produce the majority of cellular traction forces and are instrumental in rear retraction during migration [9,17,18,19]. Although these important structures have been described several decades ago, their dynamics and mechanics are far from being fully understood.

In a migrating cell, the actin cytoskeleton undergoes continuous polymerization and depolymerization as well as remodeling of the actin filament organization within existing fibers and networks. Early studies of actin dynamics in embryonic chick heart fibroblasts visualized by microinjected tetramethylrhodamine-actin indicated complex remodeling of existing fibers and bundles with multiple fusion and splitting events [20].

Later, a series of major studies more specifically targeted the origin of VSFs during cell migration. It was shown that VSF formation in osteosarcoma (U2OS) cells occurs by fiber crosslinking between stress fiber subtypes in upper cell planes. During this process, two non-contractile dorsal stress fibers connect with the ends of a contractile transverse arc, leading to formation of a VSF. When the structure resolves into a straight fiber, the focal adhesions of the dorsal stress fibers are found at both ends of the newly formed prominent VSF [13, 21]. A different scenario has been described for thinner, smaller cortical stress fibers, which can form by condensation of cortical actin mesh between two adhesions in the ventral plane of the cell [22,24]. In addition to those *de novo* VSF formation mechanisms, the number of VSFs can be increased by remodeling of the existing VSF network; when large focal adhesions undergo splitting; so that each fiber it is attached to becomes a new VSF as a result [23].

Altogether, the dynamics of VSF initiation has been fairly well studied. In contrast, VSF remodeling beyond their formation has not been analyzed in such great detail. One question left behind is how short VSFs formed close to the cell edge are transformed into long ones. Because VSFs provide forces to contract the cell body in the direction of movement, it is essential that they extend all the way along the long cell axis. Our study reported here is a step into understanding the interplay of adhesion dynamics, actin bundle remodeling and myosin contractility-driven forces, leading to VSF extension and alignment necessary for efficient cell movement.

We report a method of VSF extension in migrating fibroblasts that involves the merging of two ventral fibers at an intervening adhesion by formation of a myosin-II bridge to form a single VSF. In contrast to previously described VSF formation processes, this merging does not involve fibers from upper cell layers but occurs solely in the ventral cell plane. By experimental and computational means, we find that adhesion dynamics is a critical step for VSF merging, and that merging efficiency is guided by contractile forces exerted to the adhesion by myosin contractility. We observe that the merging process is used for dynamic reorganization of VSFs in steady state according to cellular needs such as forming protrusions, cell turning or simply to change the distribution of VSFs in the cell. In this paper we thoroughly characterize merging as a new paradigm for remodeling the ventral contractile system in cells.

## Results and discussion

### Ventral stress fibers remodel by merging at intervening focal adhesions

To analyze dynamics of ventral contractile cytoskeleton in motile MRC5 cells *(lung fibroblast cells, ATCC®),* we visualized contractile fibers and focal adhesions by expression of GFP-Myosin light chain and mCherry-Vinculin respectively. These data revealed that in process of their reorganization, VSFs consistently merge at intervening focal adhesions (***Fig 1B, Suppl Video 1***). In each merging event, a pre-existing VSF (referred to as a primary fiber) becomes involved in merging as a newly forming VSF (referred to as a secondary fiber) attaches to one of its end adhesions (referred to as an intervening adhesion).

While the primary fiber may arise by various mechanisms and exist in a cell for a significant time prior to merging, the secondary fibers predominantly form anew (***Fig 1C, Suppl Video 2)***. Visualization of GFP-Utrophin indicates that secondary fibers arise from the disordered actin mesh by spontaneous condensation of the mesh into discrete fibers. This effect is similar to cortical stress fiber formation [22], except that instead of forming with two adhesions on each end, the secondary fibers formed proceed to attach to a preexisting primary fiber for further consolidation into a VSF. Interestingly, a similar process of actin condensation between two adhesions with subsequent stress fiber fusion was described during reassembly of the actin network following complete actin depolymerization in Latrunculin B washout assays [25]. This suggests that a newly forming secondary fiber possibly involves not only condensation of preexisting actin filaments but enhanced actin polymerization at these sites.

After attachment of the secondary fiber to the intervening adhesion, the primary and secondary fibers are bridged by contractile material as visualized by myosin incorporation. In parallel, the intervening adhesion dissolves and the structure is seamlessly joined into a single, straight, contractile fiber (***Fig 1B***).

In addition to the most common scenario of merging described above, we found variants on the process (***Supplemental Figure 1s***). One variation occurs when the merging fibers pull away from the intervening adhesion (***Fig 1sA, B, Suppl Video 11***). Then, adhesion can dissolve on its own (7% of events) or at the lateral side of the merging fibers (12% of events). We reason that in such scenarios, adhesion dissolution is not likely to be a limiting factor of the merging process. The second variation involves splitting of the primary fiber along its length, so that part of the primary fiber merges with the secondary fiber and the other part exists as an independent VSF (***Fig 1sC and D, Suppl Video 12,*** 15% of events). Finally, while in the majority of events secondary fiber is newly formed, a pre-existing VSF can also play this role. In such cases (12% of events), a part of pre-existing secondary fiber attaches to the intervening adhesion and continues merging into a new VSF, while the rest of the pre-existing fiber disassembles (***Fig 1sE, F, Suppl Video 13***).

All variants of the merging process are not mutually exclusive and, in different combinations, provide a significant flexibility to VSF remodeling. In most cases, we observe that VSFs formed by merging are relatively long and thick compared to the thinner, smaller cortical stress fibers [22, our observations]. At the same time, VSF formation via merging of transverse arcs with dorsal stress fibers [13] is rarely observed in MRC5 fibroblasts. Thus, VSF merging which combines preexisting ventral fibers and actin mesh in the cell can be considered a most efficient method of large VSF formation in this cell type. We suggest that it is a significant part of contractile cytoskeleton remodeling in motile mesenchymal cells.

### Two steps of the merging process: contractile “bridge” and adhesion dissolution

Based on our observations, in order to form a uniform, merged VSF two main steps are needed: formation of the myosin-containing bridge between the primary and secondary fiber over the intervening adhesion and (2) the dissolution of the intervening adhesion. Those steps are readily detected by myosin and vinculin dynamics at the point of merging ***(Fig 2A, Suppl Video 3)***. To investigate the formation of the myosin bridge and adhesion disassembly, we analyzed myosin and vinculin intensity by linescans at the merge point over time (see representative graphs of linescans in specific frames, ***Fig 2B)***. Myosin bridge formation is manifested in the intensity of myosin at the merging point reaching the same level as myosin intensity in the initial primary fiber (***Fig 2C***, see methods for analysis details). At the same time, the maximum intensity of vinculin over the merge point decreases to cellular background levels as the adhesion disassembles ***(Fig 2C)***.

**Figure 2:**
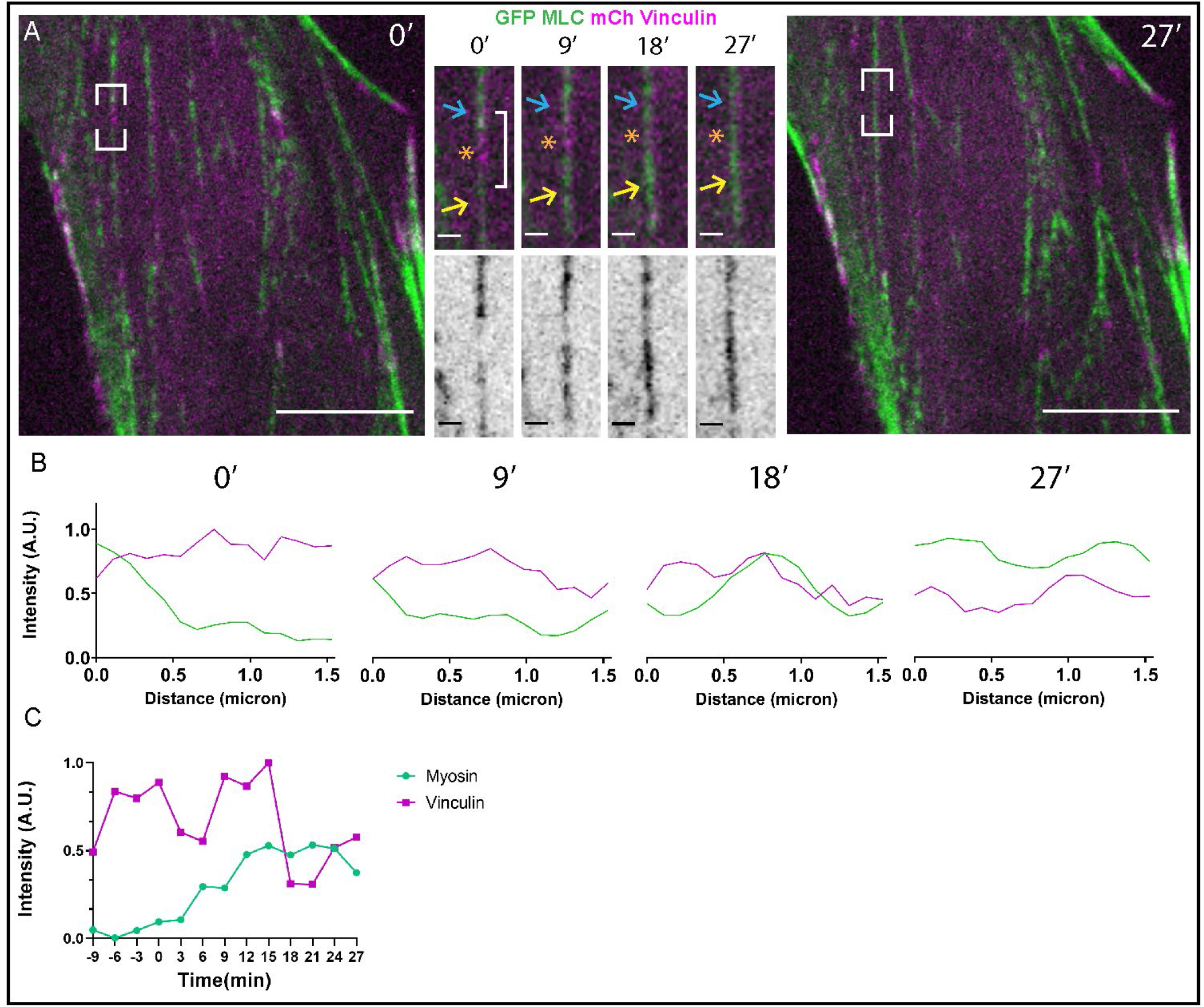
Myosin bridge formation over the intervening adhesion connects the merging fibers **A)** Ventral plane timelapse of MRC5 cell plated on 1 μg/ml Fibronectin substrate expressing GFP-Myosin Light Chain (MLC, green) and mCherry Vinculin (magenta) showing a merging event. Boxes on cell images at 0 min and 27 min note position of the merge event in the insets. Insets show the point of merging from 0 min (secondary fiber (yellow arrow) attachment to intervening adhesion (asterisk-orange above, red below) of primary fiber (blue arrow)), 9 and 18 min with merge-in-progress, 27 min with a merged fiber. Myosin bridges over the initial gap in place of the intervening adhesion as the fibers join, as seen by the continuous signal over the fiber. Scale bar= 10 μm (cell), 1 μm (insets). See Suppl Video 3. **B)** Plot profiles of Vinculin (magenta) and Myosin (green) intensity over length of the intervening adhesion at each time point. The vinculin peak at 0 min decreases as the intervening adhesion dissolves and the myosin signal increases by 27 min as the fibers are joined by the myosin bridge. **C)** Graphs of myosin and vinculin intensity parameters at the merge point over time. Myosin is denoted by ratio of minimum/maximum intensity, vinculin is denoted by maximum intensity.

These data indicate that both fiber crosslinking and adhesion dissolution are structural steps potentially important for formation of a new VSF. We propose that contractile cohesion through myosin bridging at the merge point plays a major role in forming a functionally integral fiber. In addition, the initial step to join the separate fibers could involve an actin crosslinking protein, such as alpha-actinin, which is involved in filament bundling in VSFs [26] and is enriched at focal adhesions [27,28].

Another important question is how adhesion dissolution couples into the connection of primary and secondary fibers during merging. As mechanosensitive signaling hubs, focal adhesions and actin dynamics are closely linked [4,29]. Thus, investigating how the rate of adhesion disassembly affects fiber joining is needed to understand VSF merging.

### Adhesion disassembly is needed for efficient merging

We investigated the importance of intervening adhesion disassembly in merging by plating cells on substrates with varying concentrations of fibronectin-low Fn (1 μg/ml) substrate and high Fn (20 μg/ml) substrate.

Interestingly, we found that VSF merging occurs faster on low Fn substrate (***Fig 3A, Suppl Video 4***) compared to similar event on high Fn substrate (***Fig 3B, Suppl Video 5***). Measuring the time taken to form a uniform fiber by merging showed a significant difference between low and high Fn conditions (***Fig 3C***). Investigating the dynamics at the point of merging further, we found that myosin bridge formation is also delayed in high Fn concentrations (***Fig 3D***). High fibronectin substrate concentration has been established to delay adhesion disassembly [30]. Indeed, analyzing adhesion disassembly rates in our cell model with the Focal Adhesion Analysis Server (FAAS) [31] show that rates in high Fn concentration are significantly slower than those in low FN concentration (***Fig 3E***).

**Figure 3:**
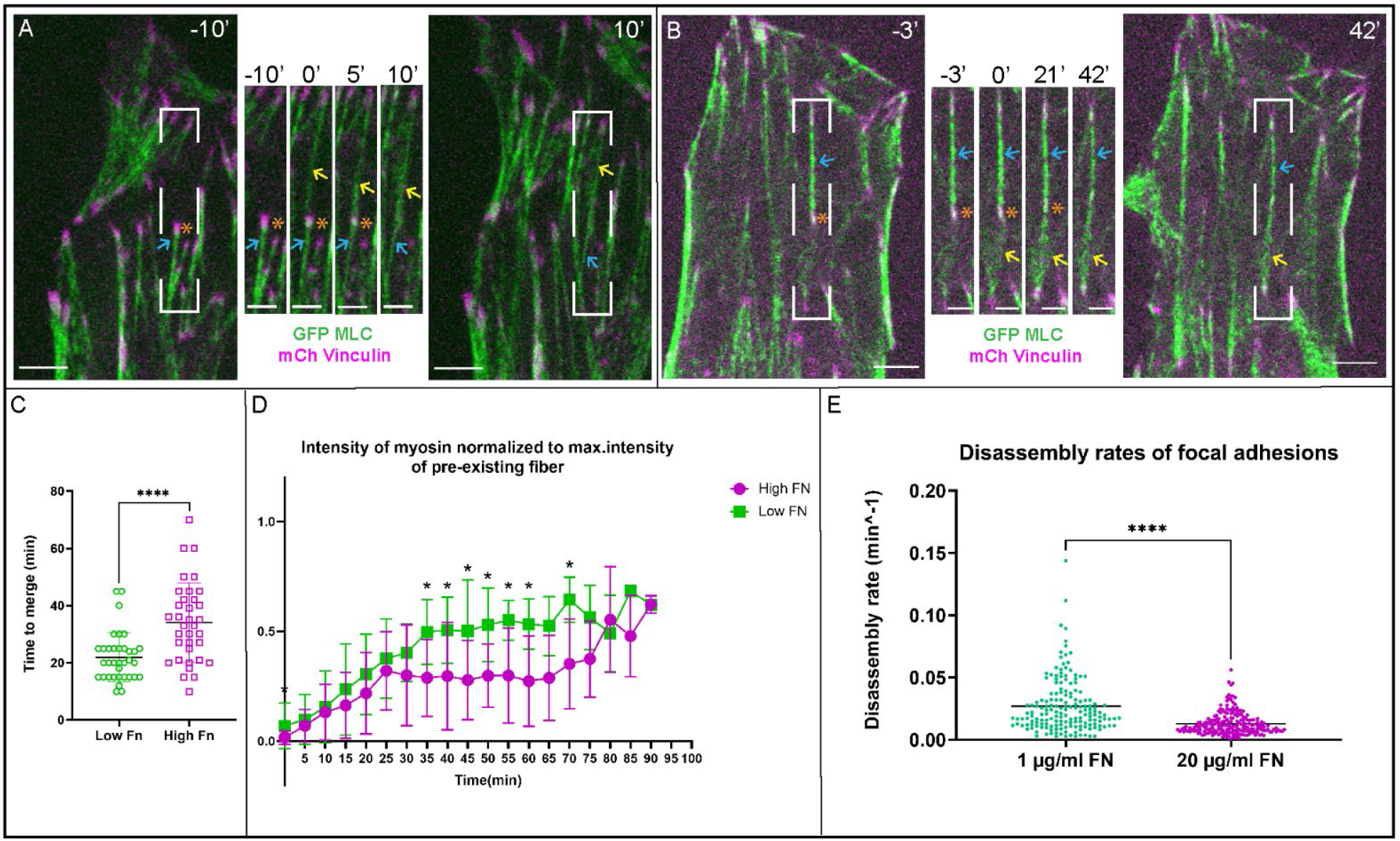
Adhesion disassembly is needed for efficient merging **A)** Ventral plane timelapse of MRC5 cell plated on 1 μg/ml Fibronectin substrate expressing GFP-Myosin Light Chain (MLC, green) and mCherry Vinculin (magenta) showing a merging event. Boxes on cell images at −10 min and 10 min note position of the merge event in the insets. Insets show merge event where a newly formed secondary fiber (yellow arrow) is attached to the primary fiber (blue arrow) at the intervening adhesion (asterisk). As the intervening adhesion dissolves, the merging fibers are joined into a single merged ventral fiber by 10 min. Scale bar= 10 μm (all). See Suppl Video 4. **B)** Ventral plane timelapse of MRC5 cell plated on 20 μg/ml Fibronectin substrate expressing GFP-Myosin Light Chain (MLC, green) and mCherry Vinculin (magenta) showing a merging event. Orange boxes on cell images at −3 min and 42 min note position of the merge event in the insets. Insets show merge event where a newly formed secondary fiber (yellow arrow) is attached to the primary fiber (blue arrow) at the intervening adhesion (asterisk). As the intervening adhesion dissolves, the merging fibers are joined into a single merged ventral fiber by 42 min. Scale bar= 10 μm (all). See Suppl Video 5. **C)** Time taken to merge (including both adhesion disassembly and continuous myosin fiber formation) for 35 events each in low and high Fibronectin conditions. Merge events on high Fibronectin substrate are significantly slower. (Mean for Low Fn= 21.97 min, Mean for High Fn= 34.06 min, Mann-Whitney test, P < 0.0001, exact, two tailed, Mean and SD on graph) **D)** Averaged graphs of myosin intensity parameters for 17 events each in low and high Fn conditions. Myosin datapoints represent the min/max intensity ratio as described in methods. Data points represent the mean and error bars represent SD. * P ≤ 0.05, T-test for normal distributions and Mann-Whitney test for non-normal distributions, two-tailed P value for either test. **E)** Graph of focal adhesion disassembly rates for cells plated in low (1 μg/ml) vs high (20 μg/ml) Fn substrate. N=172 adhesions (low Fn, mean = 0.02694 min^−1^) and 206 adhesions (high Fn, mean= 0.01299 min^−1^). Line on graph represents the mean. Mann Whitney test, two tailed, P<0.0001 (approx).

We surmise that the physical presence of the persistent adhesion in high FN conditions delays building of the contractile bridge, thus contributing to the overall delay in the merging process.

This result is not unexpected given the mechanosensitive nature of adhesion-stress fiber dynamics. We envision that when the adhesion plaque is present, tensile forces produced by both primary and secondary fibers are transmitted to the substrate (ECM), thus there is not strong force in the future bridge region. This would slow down mechanosensitive rearrangement of actin and completion of the merge. From the molecular point of view, mechanosensitive processes in effect in merging can involve not only myosin incorporation but also actin polymerization, for example via mechanosensitive mDia-1-mediated process [32,33]. Another element that could contribute to this dependency is that organization of the actin filament bundle associated with focal adhesions differs from the actin arrangement within a contractile fiber [34]. It is possible that the presence of adhesion proteins prevents a significant region of antiparallel actin filament overlap sufficient for efficient myosin incorporation.

It is important to keep in mind that adhesion dynamics also depends on tensile force applied to the adhesion by associated stress fibers [35,36,37]. The balance of forces from both primary and secondary fibers would lead to growth vs disassembly of the intervening adhesion as a consequence of merging and provide mechanical feedback between adhesion dynamics and fiber merging process.

### Smaller angle of contact at the merge point facilitates successful merging

To explore how mechanistic feedback could influence dynamics and success of merging events, we constructed a computational model of the force interactions between the primary and secondary fibers along with the intervening adhesion (***Fig 4 A,B***). Fibers are modeled as simple contractile elements while the intervening focal adhesion is modeled as a catch bond [38] whose substrate dis-association rate decreases with increasing applied force (i.e. force prolongs bond lifetime). The purpose of this modeling is to assess how the force interactions between two interacting fibers affect the dynamics of the intervening adhesion.

**Figure 4:**
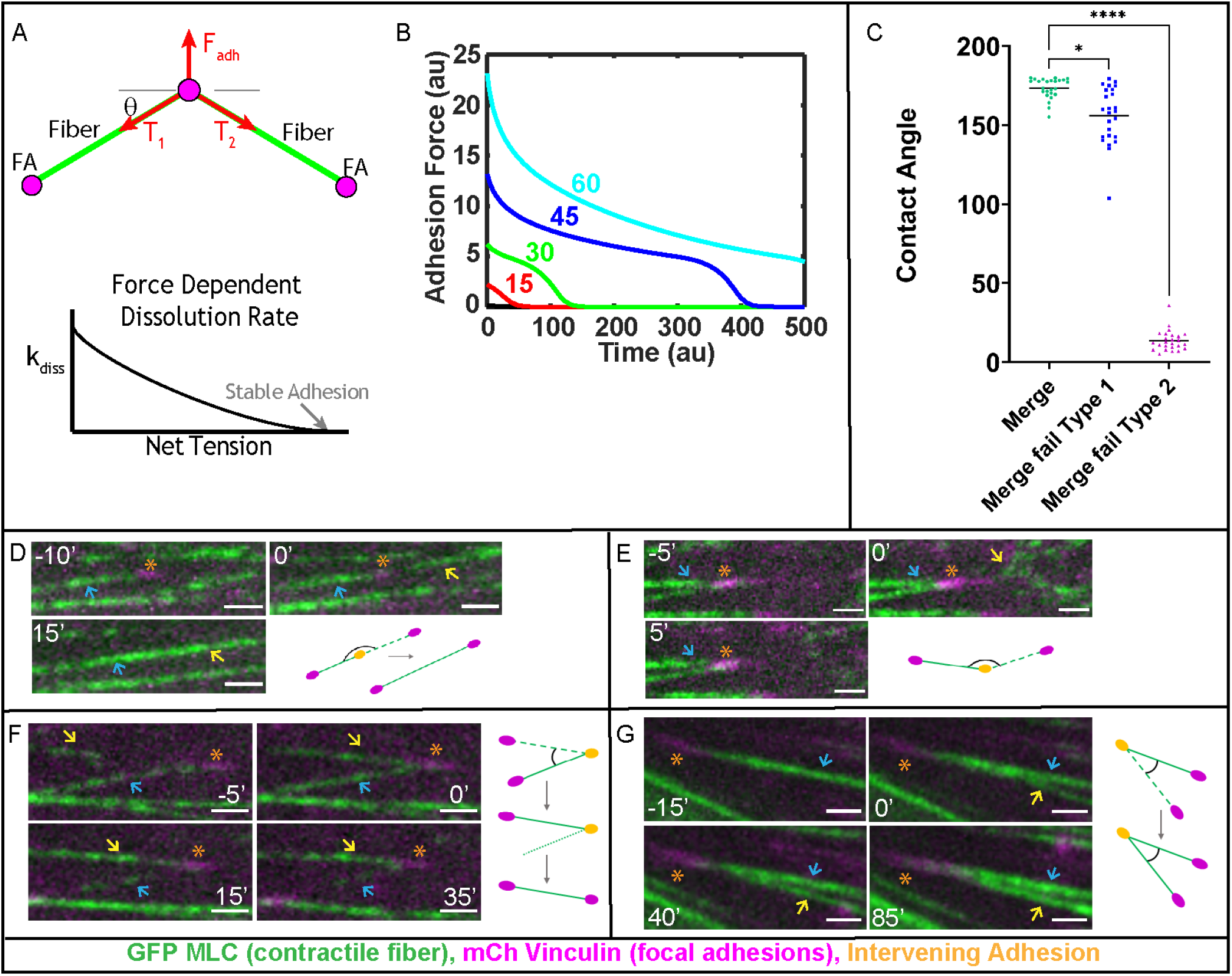
Smaller angle of contact at the merge point facilitates successful merging **A)** Top) Schematic illustration of the forces applied to an intervening adhesion between two associated stress fibers. Bottom) Schematic illustration of the force dependent stability of adhesions. The net tension of the contractile stress fibers is balanced by the adhesion-substrate force. We assume that above a certain applied force, the adhesion is stable and below this critical force the adhesion dissolves with a force dependent rate; lower forces applied to the adhesion yield more rapid dissolution. **B)** Computational simulation results showing the adhesion force as a function of the angle at which two fibers associate (θ =15,30,45,60 degrees). The slow phase of adhesion force drop results from contractile stresses straightening the joint fiber while the abrupt drop in force results from the adhesion dissolving. More acute association angles yield longer adhesion lifetimes. See Modeling Methods for further details. **C)** Graph of contact angles between the primary and secondary fiber in successful merges (Merge, mean=173.5 degrees), unsuccessful merges where the secondary fiber doesn’t connect fully (Merge fail Type 1, mean=156.1 degrees) and unsuccessful merges where the secondary fiber connects but a merging does not occur (Merge fail Type 2, mean=13.71 degrees). (n=75 total (25 each condition), Kruskal-Wallis test (two tailed), Dunn’s multiple comparisons, * P ≤ 0.05, **** P ≤ 0.0001, line represents the mean). **D-G)** Ventral plane timelapse of MRC5 cell plated on 20 μg/ml Fibronectin substrate expressing GFP-Myosin Light Chain (MLC, green) and mCherry Vinculin (magenta) showing a **D)** successful merging event **E)** unsuccessful merging event where secondary fiber doesn’t properly connect to the intervening adhesion **F)** unsuccessful merging event where secondary fiber connects, but the merge doesn’t occur due to primary fiber disassembly **G)** unsuccessful merging event where secondary fiber connects and merge doesn’t occur but V shaped structure formed remains for several minutes. In all panels, the secondary fiber is marked by a yellow arrow, the primary fiber is marked by a blue arrow and the intervening adhesion is marked by an orange asterisk. Scale bar= 10 μm. Corresponding Suppl Videos 6-9.

The main prediction of our model is that when merging fibers are positioned in a straight line, the tensile forces cancel each other, depriving the adhesion of the forces needed to stabilize it, leading to its disassembly. However, at a sharper interaction angle, the increased net force applied to the adhesion stabilizes the adhesion and at yet sharper angles, the net force exceeds the original force applied by the original fiber (***Fig 4 A,B***).

This angle of interaction dependence is a consequence of the force interactions between the two fibers and intervening adhesion. Assuming the adhesion is not accelerating, it must be in force balance such that the pulling from the two fibers is balanced by the substrate contact force. When two fibers meet at an angle such that they are nearly parallel to each other, the two contractile fiber forces counterbalance each other, leaving little net force for the substrate force to counter-balance. Due to the force dependent nature of the catch bonds, this lack of substrate force leads to dissolution of the adhesion. However, when the fibers meet at a more acute angle, the two fibers counteract each other less, leading to a larger adhesion force that stabilizes the catch bond. Thus, the angle dependent force applied to the intervening adhesion, combined with the force dependent nature of the catch bond leads to angle dependent stability of the intervening adhesion.

We have then tested this prediction by investigating how the contact angle between primary and secondary fibers in live cells would contribute to the success of merging. We found that successful merges occurred only when the initial angle between merging fibers was close to 180 degrees, such that the merging fibers are almost aligned in a straight line at the moment of initial connection (***Fig 4 B,C,D, Suppl Video 6***). In contrast, fibers that connected at an adhesion at higher angles frequently failed to merge. We have categorized failed merges based on the stability of the secondary fiber attachment to the intervening adhesion. In category 1, the secondary fiber emerges and attaches to an adhesion but exists only for a short period and dissolves before the adhesion has a chance to undergo disassembly (***Fig 4 B,C,E, Suppl Video 7***). For lower contact angles, which according to our computational model do not allow for rapid remodeling, we suggest that the attachment of secondary fibers to an existing adhesion cannot be stabilized. Instability of attachment could be explained by the alignment of actin filaments above the adhesion [34], which prevents efficient myosin incorporation. Rapid adhesion disassembly and restructuring of actin into an opposite polarity array would be needed to form a continuous contractile VSF structure, as observed in successful merges at angles close to 180 degrees (***Fig 4 B,C,D***).

In category 2, the secondary fiber attaches for an extended time, indicating that a stable connection with the adhesion has formed. In such a scenario, the failed merge is followed by either dissolution of the primary fiber (***Fig 4F, Suppl Video 8***) or the long-term persistence of both fibers in a V-shaped arrangement (***Fig 4G, Suppl Video 9***). In both cases of category 2, the angle between the two fibers is sharp, and the intervening adhesion does not dissolve in the course of observation. We speculate that in this scenario, connection of the secondary fiber to the inner side of intervening adhesion allows for rapid stabilization of attachment due to the pre-existing proper actin polarity. The merge, however, cannot be completed because the mechanosensitive feedback prevents adhesion turnover. Moreover, when a V shaped structure is formed, the adhesion often grows due to the increased pulling forces from the primary and secondary fibers, consistent with the views in the field. Overall, our analysis shows that mutual positioning of the merging fibers dictate the balance of contractile forces at the adhesion, and, consequently, the success of the VSF merging process.

### Merged fibers are aligned to the major axis of the cell

While our data indicate that VSF merging is a significant part of actin network remodeling in motile fibroblasts, we have also addressed how this process affects overall VSF network organization. In motile cells, the main contractile force must be aligned with the direction of migration, which is, in a directionally moving fibroblast, the direction of the long cell axis. Accordingly, stress fiber alignment with the long axis defines efficiency of cell translocation.

To analyze the impact of merging to the subpopulation of aligned stress fibers, we calculated the angles of merged fibers with respect to the long axis of the cell. ***Fig 5A (Suppl Video 10) and B*** shows that VSFs formed by merging are aligned within 25 degrees of the major axis of the cell. Computational modeling further shows that merging contributes to global alignment of VSFs by reducing the variance of VSF orientation in the cell (***Fig 5C, D)***. To demonstrate this, we created a computational model of the interactions between a population of stress fibers and simulated the length and angle distribution of the resulting network when they are and are not allowed to merge. When merger is present, the resulting VSF network is better aligned with the cell axis (***Fig 5D)*** and longer (5-10 μm) fibers (***Fig 5C)*** are observed. Thus, merging of VSFs both promotes the formation of longer fibers, and helps ensure those fibers are better aligned with the direction of migration. Both factors are necessary to facilitate the efficient transmission of force into directed migration.

**Figure 5:**
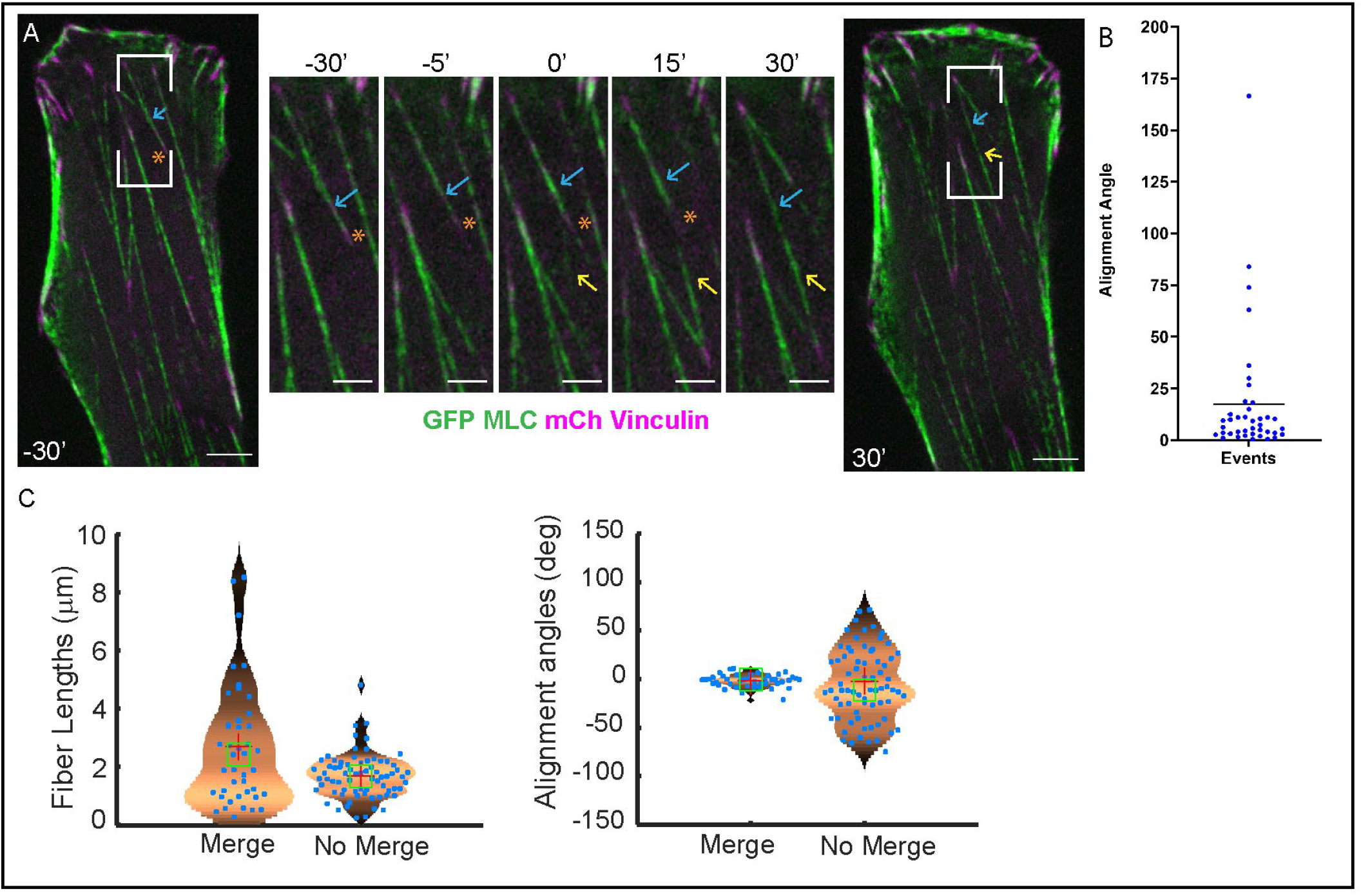
Merged fibers are aligned to the major axis of the cell. **A)** Ventral plane timelapse of MRC5 cell plated on 20 μg/ml Fibronectin substrate expressing GFP-Myosin Light Chain (MLC, green) and mCherry Vinculin (magenta) showing a merging event. Boxes on cell images at −30 min and 30 min note position of the merge event in the insets. Insets show merge event where a newly formed secondary fiber (yellow arrow) is attached to the primary fiber (blue arrow) at the intervening adhesion (asterisk). As the intervening adhesion dissolves, the angled merging fibers are joined into a single, straight merged ventral fiber that is aligned to the major axis of the cell. Scale bar= 10 μm (all). See Suppl Video 10. **B)** Graph of alignment angle of the merged fiber with the major axis of the cell (angle data points are the difference in angle between the merged fiber and the major axis) (n=40, mean=17.37 degrees, line represents the mean) **C)** Outcome of computational simulations showing the distribution of fiber lengths and alignment angles in simulations where stress fiber merger is either present or not present. Results demonstrate that when merger is present, longer fibers result and the distribution of fiber angles is less variable. See Modeling Methods for further details.

Stress fibers have been established as instrumental for cellular force production [39], and VSFs in particular are the major traction force producers in the cell [18]. Our results indicate that out of smaller stress fibers, those which are aligned similarly to each other and to the long cell axis merge more efficiently ***(Figs. 4 and 5)***. We speculate that this would enhance the VSF subpopulation coaligned with the direction of cell migration. At the same time, VSFs oriented at a bigger angle to the direction of migration cannot merge and remain small, thus only slightly contributing to the force balance in a cell. Thus, the uniform alignment of VSFs via merging contributes to directional, contractile force production by the cell, which underpins cellular functions such as cell polarization and rear-retraction during migration [11,39,40].

To conclude, in this paper, we have characterized the novel phenomenon of merging as a process for VSF formation, through experimental and computational methods. Merging occurs in the ventral plane of the cell and involves both actomyosin fiber crosslinking and focal adhesion dynamics to form a new VSF. The success of merging is dictated by the balance of contractile forces at the point of merging. In a subsequent functional feedback, merging itself directs the force landscape of the cell by modulating uniform alignment of ventral stress fibers.

## Author contribution

SN performed the cellular experiments, analyzed the data and wrote the manuscript. WRH performed computational modeling and wrote the manuscript. IK designed experimental and analytical strategy and wrote the manuscript. All authors provided intellectual input, edited and approved the manuscript.

## Materials and Methods

### Cell culture

MRC5 cells (*human lung fibroblasts, ATCC® Cat# CCL-171TM, RRID:CVCL 0440*) were maintained in MEM media (*Cat# 11095080, Thermo Fisher Scientific*) supplemented with 10% fetal bovine serum, 100 μM penicillin and 0.1 mg/ml streptomycin in 5% CO2 at 37°C. In live-cell microscopy, the cells were maintained on the microscope stage in 5% CO2 at 37°C. Cells were seeded on glass bottom dishes (*Cat# P35G-1.5-14-C, MatTek*) that were coated with fibronectin (*Cat# FC010, EMD Millipore*) 48-72 hours before experiments. Dishes were plasma cleaned before coating with 1 or 20 μg/ml fibronectin and used without plasma cleaning for coating with 10 μg/ml fibronectin.

### Expression constructs

pEGFP-gamma-Myosin Light Chain (Courtesy of Dr. Rex Chisholm, Northwestern University) was used to visualize contractile actin fibers in the cell. mCherry-Vinculin (Courtesy of Dr. Dylan Burnette, Vanderbilt University) was used to visualize focal adhesions. GFP-UtrCH (Courtesy of Dr. William Bement, University of Wisconsin-Madison) was used to visualize actin fibers and actin plaques at adhesions. Cells were transfected by the AMAXA NucleofectorTM 2b device (Cat# AAB-1001, Lonza, Program A-024) according to manufacturer’s instructions.

### Microscopy

Live-cell confocal spinning disk microscopy was used to visualize VSF dynamics. We used a CSU-X1 (Yokogawa Spinning Disk Field Scanning Confocal System) coupled with the Nikon Eclipse Ti-E inverted microscope. Images were acquired with an Apo TIRF 100× oil lens (NA 1.49) and Andor DU-897 X-11240 cameras. 3D confocal image stacks (step size: 200 nm) were captured every 1.5, 3 or 5 min, over a period of several hours. We visualized GFP-constructs with the 488 laser line, and mCherry Constructs with the 568 laser line.

### Image preparation for figures

Representative images and videos were prepared using Fiji (Ver 2.0.0 rc69-1) [41]. Maximum Intensity Projection of the ventral planes of the cell (400-600 nm total thickness) were used in representative figures and supplemental videos. In figures and videos, brightness and contrast were adjusted for each channel to make structures clearly visible (no gamma adjustments). StackReg plugin [42] in Fiji was used to align image slices through time. Representative images and videos were resized in scale though bicubic interpolation and rotated to show the relevant event.

### Data analysis

We analyzed the merging events in the cell by manually tracking the dynamics in Fiji. The point of visually observing a secondary fiber attached to the intervening adhesion and primary fiber was assigned time “0 min”. To estimate the time taken to merge (***Fig 3C***), we manually tracked the time from 0 min to the frame where both a seamless myosin bridge was present and adhesion signal was not present. 35 events each were tracked in the low (1ug/ml fibronectin substrate) and high (20ug/ml fibronectin substrate) datasets. To further quantitatively analyze the formation of the myosin bridge at the merge point over time (***Fig 3D***), we drew linescans (line width: 3 pixels) using the segmented line tool in Fiji for each time frame of a merging event. We then subtracted the background intensity and averaged the values over approx. 1 micron.

To assess myosin bridge formation, we needed a parameter indicating when the myosin intensity in the merging region becomes similar to the myosin intensity of the stress fiber. We used the ratio of minimum value/maximum value of myosin in each linescan. As the bridge formed, the ratio increased closer to 1 because of myosin incorporation forming a cohesive line between the merging fibers. If there were negative values for myosin intensity (due to signal fluctuation and background subtraction), we assigned that value to “0” (subtracted the lowest negative value in the event from each intensity value). This ensured that all min/max ratios were positive. 17 events each in low and high FN condition were averaged to form a curve in Graphpad Prism (Ver 9.0.0 for Windows, GraphPad Software, San Diego, California USA, https://www.graphpad.com).

We tested for normality of distribution using the Shapiro-Wilk test, Student’s t-test was performed for the normally distributed values, Mann-Whitney test was performed for non-normally distributed values at each time point to assess the significance.

We used the Focal Adhesion Analysis Server [31] to calculate the disassembly constants of adhesions in the low and high fibronectin substrate conditions. A semilog plot of fluorescent intensity of vinculin as a function of time was plotted by the software based on the protocol in [43]. The slope of this graph determines the apparent rate constant of adhesion dissociation. The input files were thesholded in the software according to recommended input protocol, the threshold level was chosen for each movie based on best judgement of the investigator to compensate for variable adhesion marker expression.

To calculate the angle at the merge point, we marked the angles manually at 0 min by the angle tool in Fiji. 25 events were marked for each condition in Fig 4C. Kruskal-Wallis test with Dunn’s multiple comparisons was performed to assess the significance.

To calculate the alignment of merged fibers in the cell, we fit an ellipse to the cell at the time frame of conclusion of the merge and measured the angle of the long axis. We calculated the alignment angle of the fiber as the difference between the angle of the merged fiber and the long axis angle, for 40 events in Fig 5B.

### Modeling methods

We modeled ventral stress fibers (VSFs) as contractile elements with focal adhesions (FAs) on both ends. Both VSFs and FAs possess complex internal structure and dynamics. Modeling the full scope of these dynamics is beyond the scope of this investigation. Our main purpose here is to assess 1) how merger dynamics would influence the broader network of VSFs and 2) how the dynamics of an intervening adhesion influence the merger of primary and secondary fibers. Towards this end, we model VSFs as simple elastic elements and FAs as catch bonds [38].

Each VSF is modeled as a collection of discrete nodes *{x*_*i*_*}* connected by contractile segments represented as simple Hookian springs which can compress or stretch depending on the applied forces. The first (*i=1)* and last (*i=N)* nodes represented the adhesions and the remaining internal nodes represent a computational discretization of the VSF. The internal nodes of the VSF obey the Langevin equation 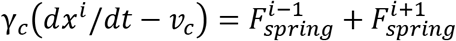. Here the term on the left is a standard drag term where the drag of the internal node is dependent on the velocity of that node relative to the speed of the cytosolic flow, which we include since the cytosol flows with the body of a moving cell while VSFs are anchored to the substrate and do not, creating a relative flow. The two spring forces account for the contractile nature of the segment connecting nodes i → i-1 and i → i+1 and take the form *F*_*spring*_ = *k*[(*x*^*i*^ – *x*^*i*+1^) − Δ*x*_*rest*_]. Here the rest length of each segment of a particular VSF is taken to be the same (i.e. a uniform discretization). The rest lengths of all segments are initialized at 0.5 μm and all fibers are initialized as strait entities such that fiber length = segment # * 0.5.

The FA nodes obey modified dynamics since they are subject to forces from the substrate as well. Force interactions between FAs and the substrate can be complex due to the catch / slip dynamics of FAs in different force regimes. Our main goal here is to simply encode the fact that FAs resist motion relative to the substrate, which we model as a drag relative to the substrate with a high drag coefficient. The i = 1 (and similar for I = N) FA node obeys the Langevin equation 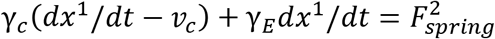. The first term is again drag imposed by the potentially flowing cytosol, the second term is drag imposed by the substrate adhesion, and the final term is the single spring force between the i=1 and i=2 nodes.

The parameters of this model do not significantly impact its dynamics. We non-dimensionalize the system such that the cytosolic drag coefficient is γ_*c*_ = 1. It may be possible to calculate the drag on a rod immersed in the cytosolic fluid. However, the precise value mainly determines the timescale over which drag forces operate in this simple model, and not the qualitative results, which are our main focus. The spring forces ensure fibers relax to a strait geometry with a determined length and the spring constant only affects the timescale of the relaxation. As such we make the choice *k*/γ_*c*_ = 1. In simulations not shown, we have taken k=0.1, 10, 100. While this spring constant can affect numerical stability (stiffer spring constant requires smaller simulation time step), this choice does not affect the qualitative outcomes presented in this article. The cytosolic flow (a consequence of cell movement) is taken to be *v*_*c*_ = 0.007 μm / s, consistent with observations of fibroblast motility speed [44].For all simulations, the computational time step is taken to be Δ*t* = 0.5 with a final simulation time of *T* = 5000.

In **Figure 4**, we model the interaction between two VSFs with these dynamics. In this specific model, one of the internal nodes represents the intervening adhesion and obeys the appropriate dynamics (cytosolic drag, substrate drag, and two spring forces). The goal of this specific model is to determine how the angle of contact influences adhesion stability. Since it is well established that, in appropriate regimes, applied forces strengthen and prevent dissolution of FAs, we consider assume that the friction coefficient *γ*E(t) is time dependent and depends on the force applied to the adhesion. This time / force dependence is modeled as *dγ*_*E*_/*dt* = −*k*_*diss*_*γ*_*E*_. The rate of dissolution is force dependent and for simplicity is modeled as a linear function where increased force yields a lower rate of dissolution: *k*_*diss*_ = 0, *F* > *F*_*max*_, and *k*_*diss*_/ *k*_*max*_ + *F*/*F*_*max*_ = 1,0 < *F* < *F*_*max*_. The *k*_*max*_ parameter controls the maximum speed of dissolution and for simplicity we set *k*_*max*_ = 1. We calibrate *F*_*max*_ so that the term inal node adh esion s are stab le a nd set *F*_*max*_ = 5. This choice does affect results somewhat. Specifically, it affects the contact angle at which adhesions become unstable. Wider ranges of contact angles become stable for smaller values of this parameter. The essential explanation for this is that this parameter defines the window of forces [0, *F*_*max*_] where the adhesion is insufficiently loaded to be stable, and more acute interaction angles produce larger loadings.

For **Figure 5**, we model the interaction between fifty initial fibers (with initial length 2.5 μm) to assess how the merger dynamics influence the length and angular distribution of resulting fibers. To create the initial condition for all simulations, we randomly insert fifty fibers allow those to merge where overlaps occur such that the new fibers are non-overlapping. We then simulate two model conditions: one where fibers that come in close proximity merge, and another where merge is not allowed. For simplicity in the merge simulations, we assume that when two fibers come into contact, they successfully merge. It is possible to incorporate rules for conditional success of VSF merger. Here however our purpose is mainly to assess how the presence of merger influences the resulting distribution of VSFs, not to model the effects of more detailed observations about mergers.

---

**Figure 1s:**
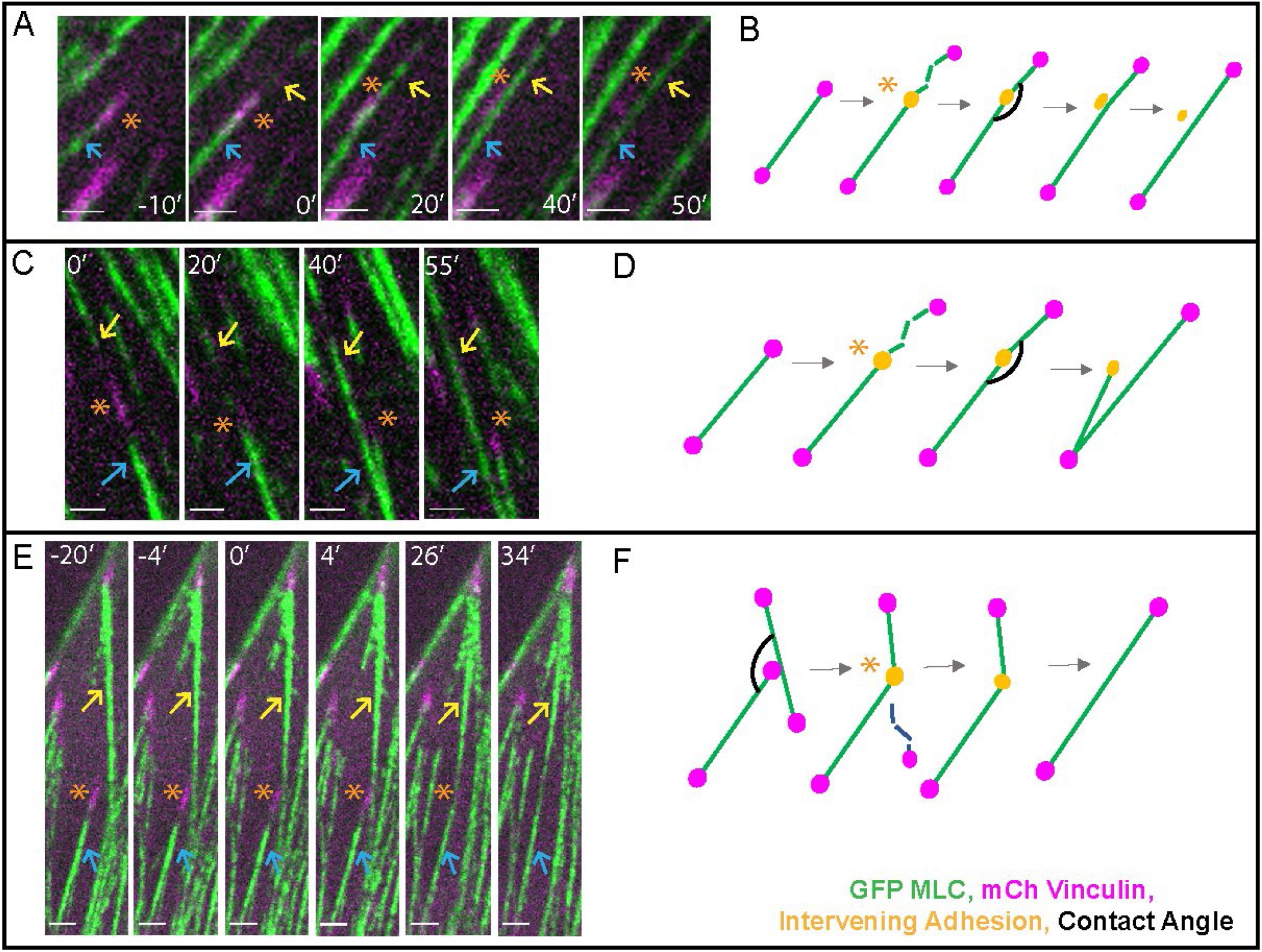
Merging steps have variants based on the nature of merging fibers and fate of the intervening adhesion. **A)** Ventral plane timelapse of MRC5 cell plated on 20 μg/ml Fibronectin substrate expressing GFP-Myosin Light Chain (MLC, green) and mCherry Vinculin (magenta) showing a merging event. The merged fiber pulls off the intervening adhesion after the myosin bridge is formed in this event. Scale bar= 10 μm. See Suppl Video 11. **B)** Schematic shows the merging process in the event in A. **C)** Ventral plane timelapse of MRC5 cell plated on 20 μg/ml Fibronectin substrate expressing GFP-Myosin Light Chain (MLC, green) and mCherry Vinculin (magenta) showing a merging event. The primary fiber and intervening adhesion split from the merged fiber after the myosin bridge is formed in this event. Scale bar= 10 μm. See Suppl Video 12. **D)** Schematic shows the merging process in the event in C. **E)** Ventral plane timelapse of MRC5 cell plated on 10 μg/ml Fibronectin substrate expressing GFP-Myosin Light Chain (MLC, green) and mCherry Vinculin (magenta) showing a merging event. The secondary fiber is a pre-existing ventral stress fiber unlike most events, where it forms anew. After attachment to the intervening adhesion, part of the secondary fiber disassembles, and the primary and secondary fibers are merged. Scale bar= 5 μm. See Suppl Video 13. **F)** Schematic shows the merging process in the event in E.

## Video Legends

Suppl Video 1: Video of VSFs merging (from Fig 1B), 5 fps, jpeg compression

Suppl Video 2: Video of secondary fibers forming by mesh condensation (from Fig 1C), 5 fps, jpeg compression

Suppl Video 3: Video of VSFs merging (from Fig 2A) showing dynamics at the merge point, 5 fps, jpeg compression

Suppl Video 4: Video of VSFs merging on low Fn substrate (from Fig 3A), 5 fps, jpeg compression

Suppl Video 5: Video of VSFs merging on high Fn substrate (from Fig 3B), 5 fps, jpeg compression

Suppl Video 6: Video of successful merge (from Fig 4D), 2 fps, no compression

Suppl Video 7: Video of merge fail type 1 (from Fig 4E), 2 fps, no compression

Suppl Video 8: Video of merge fail type 2 (from Fig 4F), 2 fps, no compression

Suppl Video 9: Video of merge fail type 2 (from Fig 4G), 5 fps, jpeg compression

Suppl Video 10: Video of VSFs merging (From Fig 5A), 5 fps, jpeg compression

Suppl Video 11: Video of VSFs merging in a variant process (Fig 1sA), 5 fps, jpeg compression

Suppl Video 12: Video of VSFs merging in a variant process (Fig 1sC), 5 fps, jpeg compression

Suppl Video 13: Video of VSFs merging in a variant process (Fig 1sE), 5 fps, jpeg compression

